# Deciphering the roles of N-glycans on collagen-platelet interactions

**DOI:** 10.1101/385179

**Authors:** Christian Toonstra, Yingwei Hu, Hui Zhang

## Abstract

Collagen is a potent agonist for platelet activation, presenting itself as a key contributor to coagulation via interactions with platelet glycoproteins. The fine-details dictating platelet-collagen interactions are poorly understood. In particular, glycosylation could be a key determinant in the platelet-collagen interaction. Here we report an affinity purification coupled to mass spectrometry-based approach to elucidate the function of N-glycans in dictating platelet-collagen interactions. By integrative proteomic and glycoproteomic analysis of collagen-platelet interactive proteins with N-glycan manipulation, we demonstrate that the interaction of platelet adhesive receptors with collagen are highly N-glycan regulated, with glycans on many receptors playing positive roles on collagen binding, with glycans on other platelet glycoproteins exhibiting inhibitory roles on the binding to collagen. Our results significantly enhance our understanding of the details of glycans influencing the platelet-collagen interaction.

---

Platelets are crucial mediators of primary hemostasis and endothelial repair, however, chronic pathological platelet activation also plays a critical role in progressive vascular occlusion diseases.^1^ Blood vessel damage exposes collagen within the subendothelial surface which serves as the primary target site of platelet action.^1^ Contact between platelet receptor proteins and nascent exposed collagen instigates platelet activation and initiates the coagulation cascade. The coagulation cascade is well defined, however, a detailed understanding of fine details dictating the platelet-collagen interaction remains elusive. Post-translational modifications, including protein glycosylation are potential key regulators of protein-protein interactions, and have been shown to contribute to platelet protein adhesion to collagen.^2 3^ The majority of critical collagen binding platelet membrane proteins, including GPIbα, GPIIb/IIIa, CD36, GPVI, and GPIX, are glycoproteins. Previous studies have reported the significant impact glycosylation on platelet-collagen adhesion for both N-glycosylation (glycoprotein VI (GPVI), integrin β1 (ITGB1)),^4 5^ and O-glycosylation (glycoprotein Ib α (GP1BA), integrin αIIb, integrin α5 (ITGA5), and GPVI).^6 7 8^ In addition, to other mechanisms, N- and O-glycosylation have been shown to play critical roles in many facets of the hemostatic system, including, expression, liver clearance ^9 10 11 12 13 14^, catalytic activity,^15^ signal transduction,^16^ as well as the proper function of von Willebrand factor (VWF) in the clotting process.^15 17 18 19 20 21^ However, there is a conspicuous dearth of information regarding the specific role of N-glycosylation dictating the binding of platelet proteins to collagen. Studies that have analyzed the impact of glycosylation on particular platelet protein function have relied predominantly on recombinant techniques for protein expression. The glycosylation of recombinant proteins varies widely based on cell type that is used to generate the proteins, and thereby limits the inferences that can be made regarding the impact of native glycosylation on the platelet protein-collagen interaction. Here we examined the roles of N-glycans on platelet protein-collagen adhesion. In this study, we developed a method to probe the specific impact of glycosylation on protein-protein interactions (PPIs) based on the interaction of immobilized “bait” with soluble “prey” glycoproteins with and without N-glycans. The approach was validated using a simple one-to-one lectin-glycoprotein proof-of-concept system and then applied to a complex collagen (bait)-platelet protein (prey) system to elucidate the roles of glycosylation in thrombus formation. This study provides insight into the mode of binding between platelet proteins and collagen.

## Results

### Proof-of-concept: *sambucus nigra* agglutinin (SNA)-fetuin binding

The schematic workflow to examine the functions of glycans in protein-protein interaction (FOGIPPI) using immobilized “bait”-soluble prey protein capture is described in Fig. 1a. As a validation of the concept of the workflow, we speculated that the known interaction between the immobilized plant-derived *Sambucus Nigra* Agglutinin (SNA), a lectin derived from elderberry bark with a known affinity for sialic acids, ^22^ and the bovine serum glycoprotein fetuin-B could provide the ideal proof-of-concept system. Fetuin-B has three N-glycosylation sites, bearing sialylated bi- and tri-antennary complex-type glycans.^23, 24, 25^ Loss of sialic acids completely abrogated SNA binding (Fig. 1b). In addition, we also observed the loss of binding of Beta-2-glycoprotein 1 to SNA after sialidase treatment (Fig. 1c). Beta-2-glycoprotein 1 is a highly sialylated bovine serum protein that is a commonly found contaminant in fetuin preparations.

**Fig. 1.**
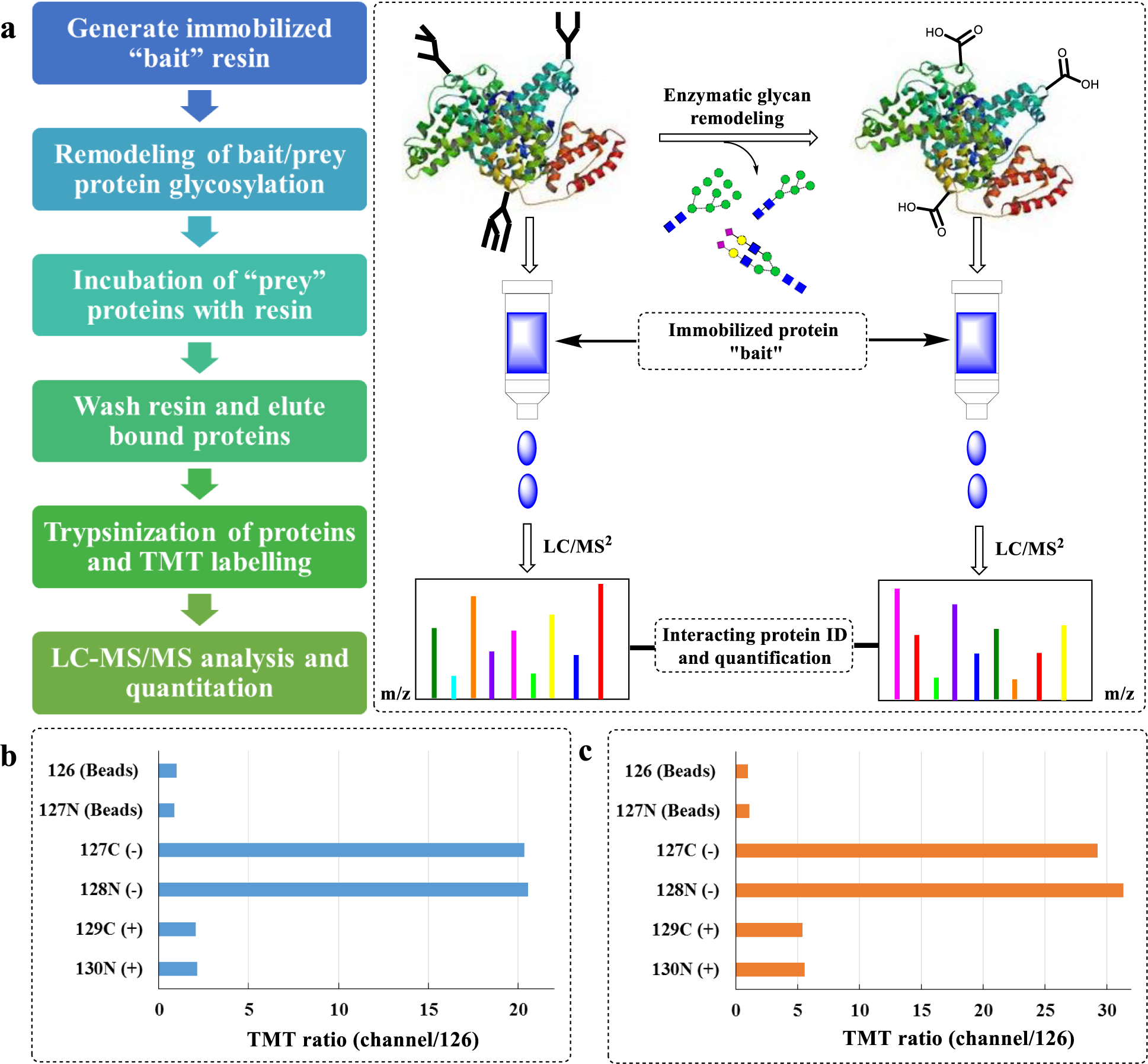
Proof-of-Concept strategy to analyze the sialic acid dependence of protein-protein interactions using a fetuin-SNA model system. **a** Schematic representing the analytical workflow. **b** N-glycan dependence of fetuin-B binding to immobilized SNA lectin analyzed via tandem mass tag (TMT) analysis, where Beads represents Tris-blocked agarose resin (reference channel), (-) represents no treatment, and (+) represents sialidase treatment. **c** Sialic acid dependence of SNA-Beta-2-glycoprotein 1 binding using TMT analysis. Each channel was quantified as a ratio compared to the blank reference channel 126.

The two forms of fetuin-B/Beta-2-glycoprotein 1 (i.e. untreated (-), and α2, 3, 6, 8- neuraminidase (sialidase) treated (+)) were incubated with SNA/blank resin. Bound proteins were eluted and digested and the trypsinized peptides were labelled with 10plex TMT reagents and subsequently analyzed via LC-MS/MS. TMT analysis largely followed expectations (Fig. 1b, c), with a drop in binding by both proteins following sialidase treatment, suggesting that alterations in protein-protein interactions due to sialic acid changes can be efficiently analyzed using the immobilized bait protein-soluble prey method. The method provides a degree of flexibility. First, the impact of bait protein glycosylation, provided the bait is a glycoprotein, can also be manipulated. Second, the changes of glycosylation on both bait and prey proteins can be achieved by glycosidases or glycotransferases.

### Determine the impact of N-glycosylation on platelet-collagen PPIs

The fine details dictating platelet receptor protein binding to collagen are critical to understanding the nature of the adhesive receptor/protein-collagen interaction, as well as the factors necessary for optimal binding. The use of quantitative proteomics provides a non-hypothesis driven characterization of PPIs between platelet proteins and collagen with altered glycosylation states. The identification of alterations in platelet protein-collagen binding following enzymatic deglycosylation will facilitate the discovery of novel, glycan-regulated protein-protein interactions with potential, downstream therapeutic applications. We applied the FOGIPPI method to platelet protein binding to collagen. Immobilized collagen resin was generated using AminoLink agarose resin. Human platelet proteins were isolated from platelet-rich plasma (PRP) and lysed in a lysis buffer containing a non-ionic detergent (Triton X-100) to ensure the solubilization of the majority of membrane-bound adhesive receptors without denaturing the protein structure or protein complexes, maintaining the capacity for protein-protein interaction (PPI). Lysed platelet proteins were treated with either PBS, PNGase F, or sialidase, and each of the treated platelet proteins were incubated with agarose-bound collagen (Supplementary Fig. 1).

Proteins were thoroughly washed and eluted under acidic conditions. Proteins released from different washing conditions were monitored by mass spectrometry analyses (Supplementary Fig. 2 and 3) to ensure binding selectivity. Acid eluted proteins were digested via trypsin, and the trypsinized platelet proteins were labelled in duplicate TMT channels and four individual collagen binding conditions were incorporated in two TMT sets (Supplementary Fig.), including native collagen and native platelet proteins (**-COL/-PLT**) [reference channel], native collagen and PNGase F-treated platelet proteins (**-COL/+PLT**), native collagen and sialidase-treated platelet proteins (**-COL/SPLT**), deglycosylated collagen and native platelet proteins (**+COL/-PLT**),and deglycosylated collagen and deglycosylated platelet proteins (**+COL/+PLT**). The negative control channel consisted of native platelet proteins and Tris-blocked agarose beads (**-/c**) (Supplementary Fig. 4). The TMT labelled peptides were pooled, fractionated using basic reverse phase liquid chromatography (bRPLC). The bRPLC fractions were concatenated (Supplementary Fig. 5) into 24 fractions and LC-MS/MS analysis was used to identify and quantify changes in collagen binding following glycan manipulation.

MS/MS analysis was operated in a data-independent manner, necessitating the use of technical replicates for each channel (Supplementary Fig. 4). Following TMT LC-MS/MS analysis of collagen-bound platelet proteins, 2,255 proteins were identified. This high protein identification number is typical of affinity enrichment followed by 2D-LC-MS/MS analysis, as proteins that bind non-specifically to the resin or indirectly to collagen through protein complex formed by collagen binding proteins are included in the analysis, indeed for many proteins, the TMT intensity values were detected in the control with the blank beads.^26^ Reproducibility analyses revealed close agreement between channels (Supplementary Fig. 6). Proteins that interacted with collagen were stratified based on both their statistical significance and binding strength, with a minimum inclusion criteria of 2.0-fold change and P value of less than 0.05 in platelet protein and collagen interactions. Volcano plots were generated in order to visualize the alterations in protein binding following glycan manipulation (Fig. 2), yielding distinct profiles. Significant alterations in protein binding following glycan manipulation was defined as ≥ 2-fold change up or down. Under such minimal inclusion criteria, 601 platelet proteins were found to have reduced binding for native collagen following PNGase F treatment of the platelet lysate, while 109 proteins were found to have reduced binding for collagen following PNGase F-treatment. Similarly, sialidase-treated platelet lysate, PNGase F-treated collagen, and PNGase F-treated collagen/platelet proteins yielded protein ratios with at least 2-fold changes (up /down) of 485/150, 209/145, and 182/179 respectively (Fig. 2). Removal of N-glycans or sialic acid from platelet glycoproteins, in general, appears to largely decrease platelet protein binding to collagen (Fig. 2a, 2b), indicating an overall inhibitory role for N-linked glycans or sialic acids of platelet glycoproteins in platelet protein-collagen interactions. Conversely, collagen deglycosylation generally appears to result in higher protein binding (Fig. 2c, d), indicating that the glycans on collagen may play some negative role in platelet protein-collagen interactions that limits binding.

**Fig. 2.**
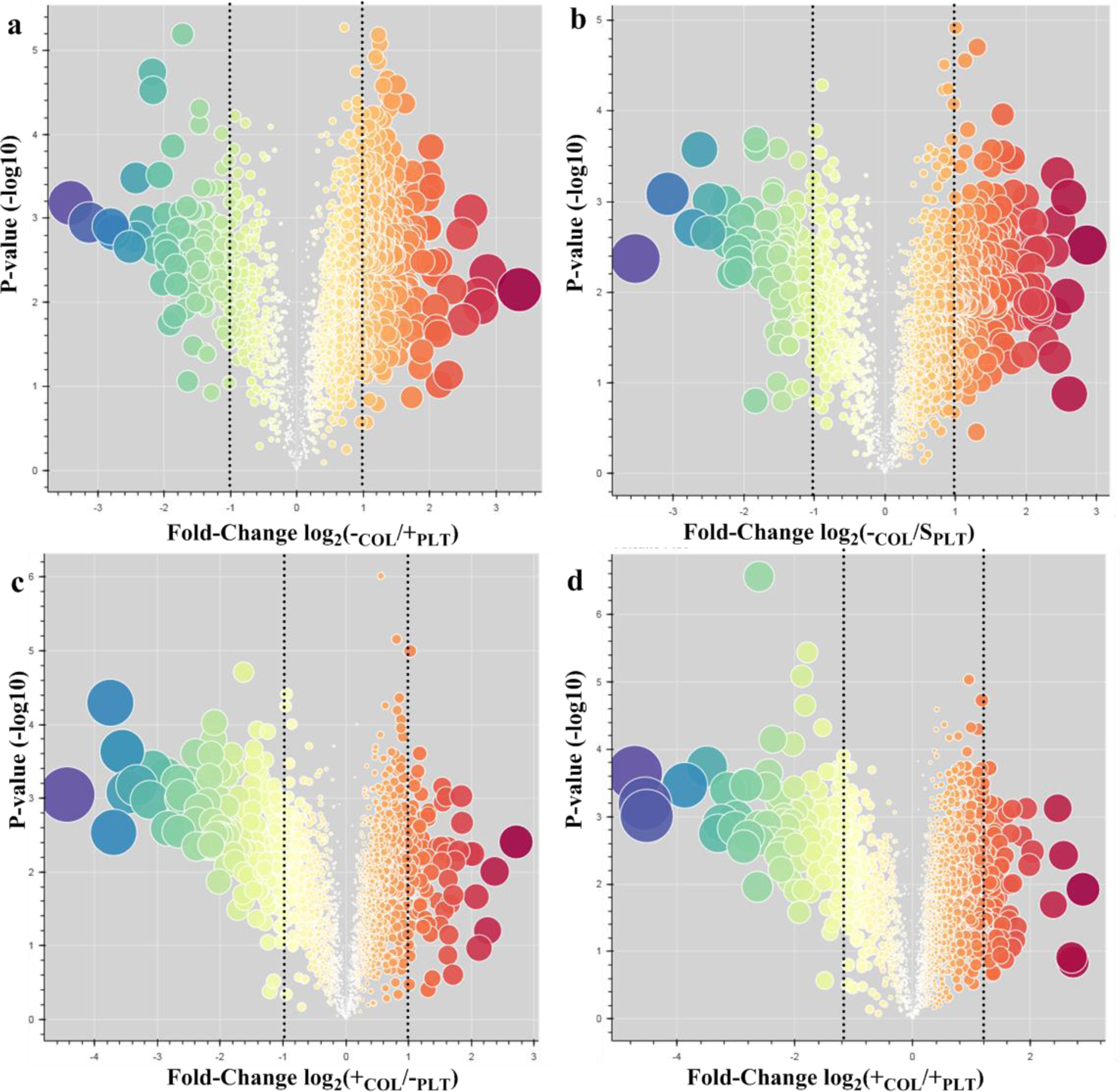
Comparative analysis of the loss of collagen-binding affinity between platelet proteins (PLT) and immobilized collagen (COL) following subsequent PNGase F (+) or sialidase (S) treatment of either PLT or immobilized COL via TMT-labelled quantitative proteomics. Volcano plots of all quantified proteins displays the relationship between statistical significance and fold-change. The log2 fold-change (*x axis*) was plotted against the –log10 p-value (y-axis). 2-fold changes and P value of less than 0.05, indicating a loss or gain of binding, occur to the right or left, respectively, of the dashed lines. **a.** Binding of PNGase F-treated PLT to untreated collagen (-COL/+PLT). **b.** Binding of sialidase-treated platelet proteins to untreated collagen (-COL/SPLT). **c.** Binding of untreated platelet proteins to PNGase F-treated collagen (+COL/-PLT). **d.** Binding of PNGase F-treated platelet proteins to PNGase F-treated collagen (+COL/+PLT).

### Intact glycopeptide analysis identifies the specific glycoproteins and their glycan changes associated with the alterations in PPI

In order to rationalize the N-glycan dependency of binding interaction between platelet adhesive receptors/proteins and collagen, we analyzed the platelet sample using intact glycopeptide analysis. Intact glycopeptide analysis allowed us to site-specifically analyze individual glycans from a complex mixture of proteins. In order to estimate the contribution of intact glycopeptides from platelet versus contaminating plasma, samples were prepared from either washed platelets or platelet-free plasma. Abundant serum proteins were removed using a multiple affinity removal system-6 (MARS-6) (Fig. 3a). The intact glycopeptide data was searched using the software package GPQuest 3.0.^27^ In total, 3111 unique glycopeptides were identified with a 1% FDR for the platelet sample, corresponding to 192 distinct N-glycans from 278 glycoproteins. For the plasma sample, 3452 unique glycopeptides were identified (1% FDR), including 216 distinct N-glycans, corresponding to 181 glycoproteins. Analysis of the overlapping glycopeptides shared between platelet and plasma, indicated a moderate number of glycopeptides (819) in common between the two samples (Fig. 3b). The majority of glycopeptides identified between the two samples were unshared, however, there was significant overlap at the glycoprotein level, with 40% of the glycoproteins shared between platelet and plasma, with only 15% of the glycoproteins identified unique to plasma (Fig. 3c). The disparity between the identifications at the glycopeptide versus glycoprotein levels, suggests that the patterns of glycosylation between platelet and plasma samples is different. While the peptide portion of a given protein may be shared, the glycan portion appears to be largely unique and source specific. The unique N-glycans identified from platelet and plasma intact glycopeptide samples were divided into the three major categories (i.e. high-mannose, hybrid-type, and complex-type), and demonstrated a very similar distribution (Supplementary Fig. 8a, b). Intact platelet glycopeptides appeared to have slightly higher levels of oligomannose glycans, while intact glycopeptides from plasma were enriched in sialic acid, and to a lesser extent fucose (Supplementary Fig. 8c, d).

**Figure 3.**
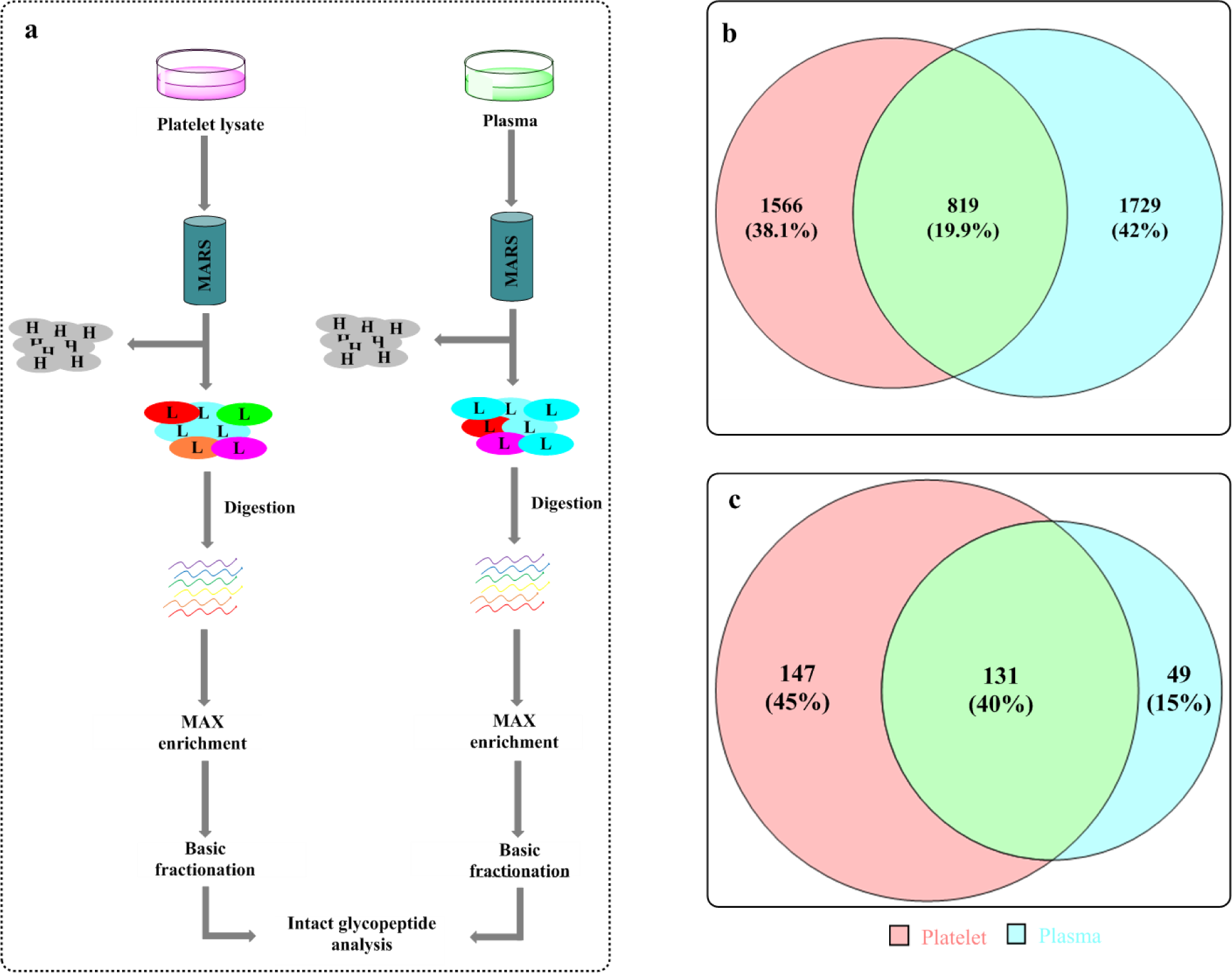
Intact glycopeptide analysis of platelet and plasma glycoproteins. **A.** Schematic illustrating the preparation of glycopeptides for intact glycopeptide analysis of platelet proteins after MARS-6 depletion of highly abundant plasma proteins. Glycopeptides were separated from non-glycosylated peptides using MAX enrichment. Sample complexity was reduced via bRPLC prior to intact glycopeptide analysis. **b.** Overlap of identified glycopeptides between platelet and plasma samples. **c.** Overlap of identified glycoproteins between platelet and plasma samples.

In addition, the *Equus caballus*-derived collagen “bait” protein that was used to generate the immobilized collagen affinity column was analyzed by LC-MS/MS. Intact glycopeptide analysis indicated that the starting collagen was a heterogeneous mixture of proteins, composed primarily of type VI, type I, type XII collagen. A number of lower abundant serum and ECM proteins were also identified, including the coagulation-related proteins fibromodulin and fibronectin. Both type VI and XII collagens are known to contain multiple N-glycans, while type I collagen has a single N-glycosylation site within the proprotein region alone. Presumably type VI and XII are the major contributors to the observed changes in PPIs following PNGase F-treatment of the collagen given the extensive N-glycosylation of both proteins. The N-glycosylation profiles of collagen were analyzed using intact glycopeptide analysis, indicating primarily highly processed complex-type glycans (Supplementary Table 1).

Collagen-enriched proteins in the TMT data set shared 10% unique glycoproteins with the platelet-derived sample, with only a 2.8% overlap with unique plasma-derived glycoproteins (Supplementary Fig. 7). The results suggest that the majority of the glycoproteins binding to collagen were from platelets, rather than contaminating plasma glycoproteins. Based on intact glycopeptide analysis, the majority of platelet-derived glycopeptides identified lacked significant sialylation in comparison to plasma proteins. The observation is not unexpected as platelets are known to be enriched with sialidase enzymes that are secreted and remodel the glycosylation patterns of the platelet glycoproteins, with desialylation accelerating platelet clearance.^13 10^

### Major platelet adhesive receptors and proteins require N-glycans for collagen binding

We stratified the identified proteins that were significantly impacted by de-glycosylation based on interaction and biological relevance. In particular, extracellular matrix (ECM)-binding proteins, glycan binding proteins, and coagulation-related proteins were extracted from the data to look in detail at the role of N-glycosylation in platelet protein-collagen interactions within the coagulation cascade. We identified 12 collagen binding proteins, six platelet glycoprotein receptors, nine integrin proteins, five fibronectin binding proteins, 11 ECM binding proteins, eight glycan-binding proteins, and 20 coagulation-related proteins. We visualized the proteins of interest via the generation of heatmaps (Fig. 4a-g). Many of the proteins highlighted exhibit pronounced alterations in platelet protein and collagen interactions following de-glycosylation of the N-linked glycans or sialidase-treatment on either the platelet proteins or the immobilized collagen. In particular, the major platelet adhesive receptors exhibited a strong dependence on platelet N-glycosylation, with a significant loss of binding following de-N-glycosylation (Fig. 5d, f).

**Figure 4.**
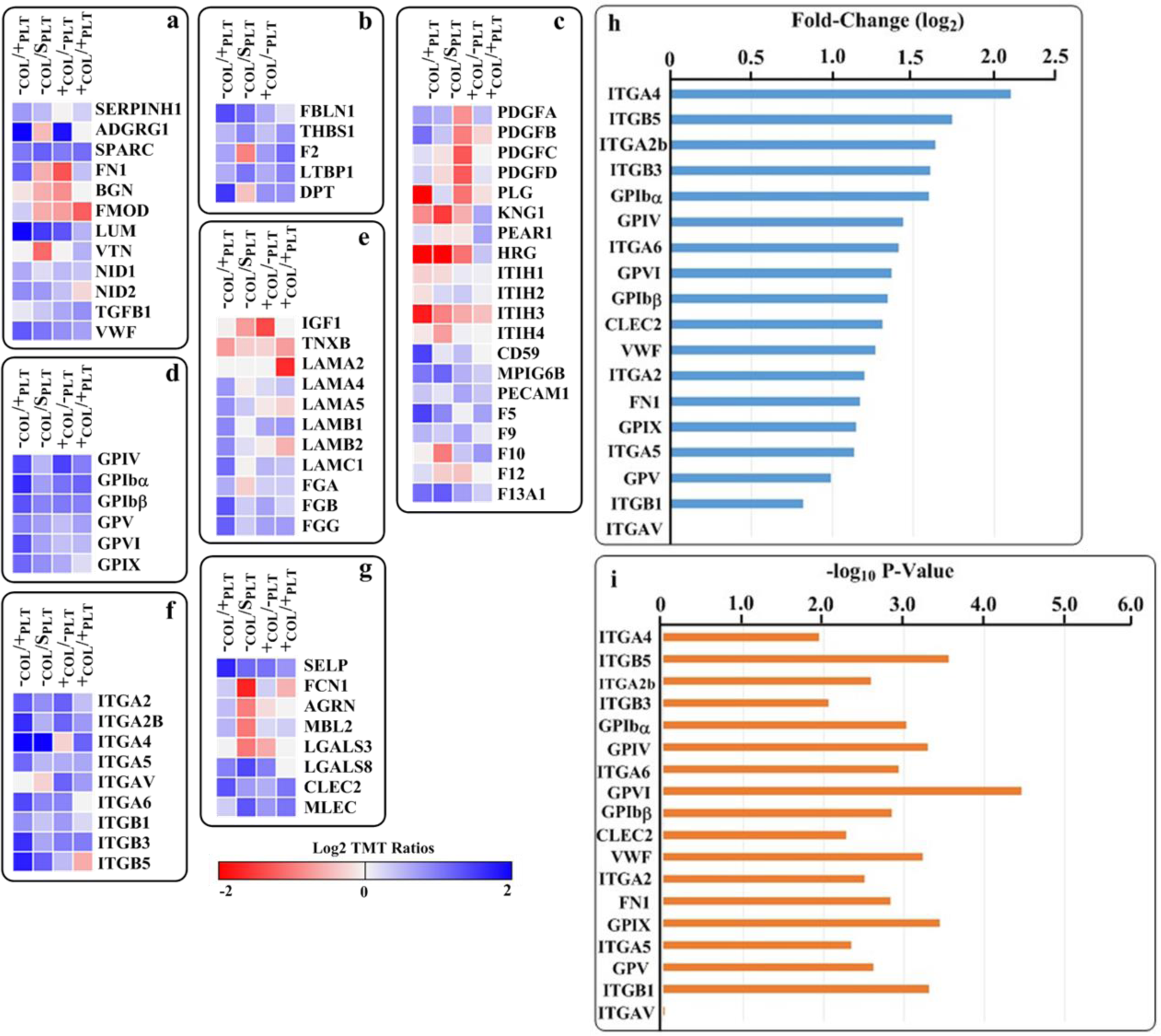
Glycan dependence of platelet protein-collagen binding for coagulation-related platelet proteins. Analysis of collagen bound adhesion receptors and other coagulation-related proteins. For each protein, the average log2 TMT ratio was calculated by dividing the untreated reference channel (-COL/-PLT) versus the PNGase F/sialidase treated channels (+/S). **a.** Heat map of identified collagen-binding proteins. **b.** Fibronectin-binding proteins. **c.** Coagulation-related proteins. **d.** Platelet glycoprotein receptors. **e** ECM-binding proteins. **f** Platelet integrins. **g** Glycan-binding proteins. **h.** Fold-change (log2 TMT ratio) of major platelet adhesive receptors/proteins indicates a strong N-glycan dependence for most proteins. **i.** P-values (-log10) of identified major platelet adhesive receptors/proteins.

To further assess the role of N-glycosylation on the adhesive interaction between platelets and collagen, we looked in detail at the 18 major adhesive proteins and receptors identified in this study, and compared the TMT ratios of the fully glycosylated platelet proteins versus the de-glycosylated platelet proteins, finding a significant (i.e. ≥ 2-fold) loss of binding following PNGase F treatment for 16 of the 18 proteins identified (Fig. 5e). The adhesive receptors/proteins can be broadly divided into two categories, the first are collagen binding adhesive receptors/proteins and the second are other ECM binding proteins. The first categories can be subdivided in to two classes, including proteins that bind directly to collagen, and proteins that bind indirectly to collagen via collagen-binding proteins. The second category includes proteins that bind to other constituents of the ECM beyond collagen, including podoplanin, thrombospondin, and laminin.

Among the identified adhesive receptors/proteins that bind directly to collagen, both of the major collagen binding receptors, GPVI and (ITGA2/ITGB1), were identified and both GPVI and ITGA2 demonstrated a > 2-fold loss of binding following PNGase F treatment of the platelet lysate, while ITGB1 was more moderately impacted (Fig. 5h). Neither receptor was significantly impacted by either sialidase treatment or collagen glycosylation. The findings for GPVI were in agreement with a prior publication which found that PNGase F-treated recombinant, GPVI, had a lower affinity for collagen.^28^ Von Willebrand Factor (VWF), a central collagen binding protein, and the first adhesive protein to initiate platelet tethering to collagen ^29^ was found to be critically dependent upon both N-glycosylation and sialylation (Fig. 5a, e) but largely unaffected by collagen glycosylation. Previous reports have shown that N-glycosylation negatively impacts platelet adhesion to VWF in the absence of collagen,^30^ however, our data indicates a positive role for N-glycans in VWF adhesion to collagen, at least under non-shear conditions. Consistent with previous studies,^31 32^ we found FN1-collagen binding was positively influenced by N-glycosylation, and binding by FN1 to collagen dropped more than 2-fold following PNGase F treatment. In agreement with previous studies,^33 34^ we found the PPI between vitronectin (VTN) and collagen to be enhanced 2.2-fold following sialidase treatment (Fig. 5a).

Many of the identified platelet adhesion receptors bind indirectly to collagen through ECM binding proteins, including VWF, FN1, and VTN. One of the most important platelet receptors is the GPIbα, GPIbβ, GPV, and GPIX (GPIb-V-IX) receptor complex that forms the incipient platelet tether to VWF bound to the nascently exposed collagen. GPIb-V-IX binds to the high-affinity surface created when VWF unrolls from a globular to a filamental conformation on the exposed collagen surface at the site of vascular injury.^35 36^ Each individual component of the GPIb-V-IX receptor complex demonstrated a concomitant loss (i.e. > 2-fold) of binding presumably following the loss VWF binding to collagen, and, with the exception of GPIbα/β, was similarly unaffected by collagen glycosylation (Fig. 5h). Similarly, FN1 plays a central role in platelet adhesion, and acts as the substrate for integrins ITGA2b/ITGB3, ITGAV/ITGB3, ITGA4/ITGB1, and ITGA5/ITGB1.^37^ With the exception of ITGAV and ITGB1, the remaining FN1 adhering integrins identified exhibited a ≥ 2-fold loss in binding following PNGase F treatment of the platelet lysate (Fig. 5h). The FN1/VTN binding receptor integrin α_v_β_3_ was differentially impacted by platelet glycan manipulation, with ITGAV exhibiting little change in binding, while ITGB3 binding was found to be highly dependent upon N-glycosylation, demonstrating a 3-fold drop in binding following PNGase F treatment of the platelet lysate. The finding was in agreement with previous literature.^38 39 40^

A number of ECM-binding platelet adhesive receptors were identified that targeted constituents of the ECM other than collagen. In the present study, glycoprotein IV (CD36), a multifunctional receptor known to bind to thrombospondin, was identified and found to be N-glycan dependent, demonstrating a sensitivity to PNGase F treatment of the platelet lysate and collagen (Fig. 5b). In addition, several receptors bind to alternative ECM proteins, such as laminins which are concomitantly exposed with collagen during vascular injury. ITGA6/ITGB1, which is a known laminin-binding receptor, was shown to lose affinity following de-glycosylation of N-glycans from the platelet proteins (Fig. 5h). The glycan binding protein C-type lectin-like receptor-2 (CLEC-2), which serves a critical role in platelet activation, ^41^ was also found to have lower affinity for the collagen resin following PNGase F treatment of the platelet lysate. The natural agonist for CLEC-2, podoplanin was not identified in this study, however, CLEC-2 exhibited a 2.5-fold decrease in binding following de-N-glycosylation. For the majority of platelet adhesive receptors/proteins that demonstrated a large binding change, the p-values were calculated and found to be statistically significant (Fig. 5i).

### Roles of glycans on platelet protein-collagen binding

Following the observation that the binding of many adhesive receptors and proteins to collagen was highly dependent on N-glycosylation, we analyzed the N-glycopeptides from the platelet adhesion receptors and collagen binding proteins that demonstrated a ≥ 2-fold change (positive or negative) in collagen binding following PNGase F treatment. Using data generated from our intact glycopeptide library (Fig. 3), we identified glycopeptides from five collagen binding proteins and nine platelet receptor proteins that exhibited reduced/enhanced collagen binding following PNGase F/sialidase treatment of platelet lysate (Table 1). Many of the glycopeptides that were identified corresponded to major collagen-binding receptors and proteins. While glycosylation of the soluble adhesive proteins exhibited the expected abundance of complex and hybrid-type N-glycans (Table 1a), many of the adhesive receptors (i.e. integrins and platelet glycoproteins) were found to contain an abundance of oligomannose glycans (Table 1b). The observation was surprising, considering the low degree of oligomannose glycans commonly identified on mammalian plasma proteins.^42^ However, a number of megakaryocyte receptors, including ITGB3 have been shown to bear underprocessed N-glycans even after receptor maturation.^43 44^ The abundance of oligomannose glycans on platelet receptor proteins suggests a structural rather than an electrostatic function for N-glycosylation of platelet receptor proteins, as no significant binding change was observed following sialidase treatment (with the exception of ITGA4). Indeed, an indirect structural role for N-glycosylation, rather than a direct glycan function has been recently suggested for ITGA2b/ITGB3 and ITGAV/ITGB3 ligand binding.^40^ As a counter example, the ECM binding protein, histidine-rich glycoprotein (HRG), demonstrated increased binding to collagen following either PNGase F or sialidase treatment (i.e. ~5.0 and ~4.0-fold, respectively). The binding behavior of HRG could be rationalized based on HRG’s substrate binding mode. HRG is known to bind to a number of negatively charged substrates via a cationic histidine-rich region.^45^ Using intact glycopeptide analysis of the platelet lysate, we identified 26 unique intact glycopeptides from HRG, all of which contained highly processed hybrid and complex-type glycans, containing 23% sialylation. The loss of sialic acid via either PNGase F or sialidase treatment reduces charge repulsion with the HRG binding substrate and presumably results in improved binding affinity. The only adhesive receptor that demonstrated a significant dependence upon electrostatic interactions was ITGA4, which was more significantly impacted by sialidase treatment that by treatment with PNGase F. It should be noted that sialidase treatment, unlike PNGase F, is not specific for N-glycans, and could be related to changes to O-glycans as well.

**Table 1.** Tabulated intact glycopeptide analysis results. **a** Intact glycopeptide analysis of collagen-binding proteins reveals primarily complex/hybrid-type glycosylation. **b** Intact glycopeptide analysis of platelet integrin and glycoprotein receptors reveals an unusually high degree of oligomannose glycans compared to serum proteins. *Unless otherwise indicated, the highest fold-change corresponded to PNGase F treatment of platelet lysate (i.e. –COL/+PLT).

|  |  |  |  |  |  |
| --- | --- | --- | --- | --- | --- |
| **a** | **Protein** | **# Glycopeptides** | **Highest Fold-Change\*** | **% Sialylation** | **% Oligomannose** |
|  | LUM | 3 | 3.9 | 0 | 0 |
|  | SPARC | 10 | 3.1 (-COL/S<sub>PLT</sub>) | 0 | 10 |
|  | VWF | 19 | 2.4 | 23.5 | 10.5 |
|  | FN1 | 29 | 2.3 | 22.2 | 6.9 |
|  | VTN | 24 | 0.46 (-COL/S<sub>PLT</sub>) | 22.7 | 8.3 |
| **b** | **Protein** | **# Glycopeptides** | **Highest Fold-Change\*** | **% Sialylation** | **% Oligomannose** |
|  | ITGA2 | 44 | 2.3 | 36 | 43.2 |
|  | ITGA2B | 4 | 3.1 | 25 | 25 |
|  | ITGA5 | 7 | 2.2 | 0 | 71.4 |
|  | ITGA6 | 54 | 2.7 | 32.7 | 9.3 |
|  | ITGB3 | 136 | 3 | 28 | 39.7 |
|  | ITGB5 | 11 | 3.3 | 0 | 81.8 |
| | GPIbα | 9 | 3 | 77.8 | 0 |
|  | GPIV | 49 | 2.7 | 39.4 | 32.7 |
|  | GPV | 18 | 2 | 11.1 | 0 |

## Discussion

A number of landmark studies using AP-MS have established methods to study the impact of PTMs on PPIs.^46 47 48 49 50 51 52^ Established AP-MS methods are hampered by their reliance on specific mAbs and/or the use of expression tags to isolate interacting species. Here we developed a TMT-based AP-MS method to identify the specific impact of N-glycosylation on the function of platelet adhesive receptors/proteins interacting with collagen, using human-derived, non-recombinant platelet proteins. The described method overcomes many limitations associated with traditional AP-MS, and provides unprecedented access to probe the function of N-glycans in PPIs. Our results demonstrate that quantitative interaction proteomics using a single immobilized “bait” protein and multiple soluble “prey” proteins provides a valuable non-hypothesis driven approach to analyze the N-glycan dependence of PPIs. The use of quantitative MS provides direct information on both the identities of prey proteins as well as the strength of the interaction based upon multidimensional perturbations in the system (i.e. glycan loss from the bait/prey).

Platelet adhesion to collagen is a multistep process requiring the sequential interaction of multiple adhesive receptors (GPIb-IX-V, GPVI, and Integrin α_2_/β_1_) with the extracellular matrix components collagen/VWF to facilitate secure tethering of the platelet to the disrupted vascular endothelium.^29^ Adhesive proteins participate in dual roles in the coagulation process by both activating platelet receptors, as well as bolstering the adhesions by acting as “cement.” Many of the major adhesive proteins are also known glycoproteins.^29^ The application of affinity purification coupled to mass spectrometry (AP-MS) to elucidate the function of glycosylation in protein-protein interactions (FOGIPPI) provides a unique and powerful method that can potentially identify glycans that regulate coagulation-related proteins. Expanding our understanding of the role of glycosylation in platelet-collagen interactions has the potential to uncover novel therapeutic pathways.

The approach was validated using an SNA-fetuin-B proof-of-concept system demonstrating the feasibility of the immobilized “bait”-soluble “prey” concept to study the function of glycans in PPIs. Application of the method to a complex system composed of immobilized collagen and platelet lysate revealed a striking glycan dependence among primary collagen-binding proteins. Using the immobilized “bait’, soluble “prey” method, we generated the first extensive glycan-dependent interaction analysis, identifying all major primary platelet adhesive receptors/proteins and revealing that the majority, including GP6, GPIb-V-IX, and integrin α_2_β_1_, are strongly reliant upon N-glycosylation for proper binding interactions. It is likely that N-glycosylation could impact substrate binding either by direct facilitating or interfering protein-protein interactions, or by modulating the optimal orientation of the glycoproteins to facilitate PPIs. The use of intact glycopeptide analysis of the platelet lysate assisted in rationalizing the structural role of N-glycosylation in the PPIs, although direct assignment of glycan function cannot be determined from this method. Intact glycopeptide analysis revealed the preponderance of oligomannose glycans, particularly among integrins, suggesting a structural rather than electrostatic role for glycans, as would be expected from more highly processed, sialylated glycans (Supplementary Table 2).

Despite the advantages and utility afforded by the immobilized “bait”/soluble “prey” method for determining the role of N-glycosylation in PPIs, there are a number of limitations that should be considered. First, endemic to all AP-MS experiments is the presence of non-specific binding proteins in addition to the proteins of interest. In our analysis, a number of non-glycosylated cytoskeletal proteins were impacted by N-glycan manipulation, however the change in affinity was likely an artifact of the alteration of charge/hydrophobicity following deglycosylation/desialylation. The changes of these protein binding to collagen could also be the result of their binding to the primary proteins whose binding to collagen is regulated by glycosylation. The issue of co-purifying non-specific binding proteins could potentially be addressed by incorporating unspecific binders as crucial elements in the bioinformatics pathway to determine the background binding as described in this study using conjugated resin as control, the method was described in other AP-MS studies such as the recent study in a LFQ/SILAC AP-MS yeast protein study.^26^ Similarly, glycoproteins that bound more strongly to the bait following deglycosylation may have bound non-specifically via hydrophobic/van der Waals forces, rather than an increased specific binding affinity; accurately determining true interactions versus non-specific interactions remains a challenge for this method. In addition, this method is limited to identifying the impact of N-glycosylation on known PPIs by analyzing the N-glycosylation changes after removing N-glycans. Novel glycan dependent PPIs cannot be confidently assessed, as the alteration in binding could arise from the generation of a non-natural binding site created by the loss of a glycan, rather than a conformational change at a biologically relevant binding site. Finally, the FOGIPPI method identifies both direct and indirect interactions. The identification of indirect interactions expands the utility of the method by allowing us to probe the impact of N-glycosylation on the majority of the components of the platelet adhesion process, however, it also significantly increases the complexity of the data analysis. Moreover, some receptors which bind to collagen-binding proteins may exhibit a change in binding due either to the loss of N-glycosylation, or the loss of agonist binding to collagen, or both. Determining the source of the loss/gain of affinity by indirectly binding proteins with the immobilized “bait” is difficult to determine with confidence.

N-glycosylation appears to play a major role in platelet receptor binding to collagen and other adhesive proteins. However, given the limited impact of sialidase treatment (-COL/SPLT) on platelet receptor proteins, coupled with the abundance of oligomannose glycans identified during intact glycopeptide analysis, we speculate that N-glycosylation may play a largely structural role in defining the binding interactions. The glycans may be required to maintain an optimal conformation that is critical for binding, and the loss of the glycan may impact binding. In addition, the abundance of oligomannose glycans on platelet receptor proteins, is interesting and suggests an attenuated N-glycan maturation pathway at the megakaryocyte level.

In summary, we have developed an FOGIPPI method utilizing proteomic approach to determine the protein-protein interactions to probe the specific function of N-glycosylation in PPIs. We have applied the method elucidate the interaction between platelet adhesive receptors/proteins and collagen, finding that the majority of glycoproteins critical for the coagulation pathway are highly N-glycan dependent for proper adhesive function. Our first systematic FOGIPPI approach provides a rich resource to study the global impact of N-glycosylation on PPIs within the context of a complex, non-recombinant system.

## Methods

### General

Platelet concentrates were obtained from ZenBio (Research Triangle Park, NC, USA). Collagen I was purchased from ChronoLog (Havertown, PA, USA). Amine-reactive resin (AminoLink) was purchased from Thermo Scientific (Rockville, IL, USA). Unless otherwise indicated, all reagents were obtained from Sigma-Aldrich (St. Louis, MO, USA).

### Platelet lysate preparation

Fresh platelet rich plasma (PRP) in acid citrate dextrose buffer was diluted 1:1 (v/v) in HEPES buffer (140 mM NaCl, 2.7 mM KCl, 3.8 mM HEPES, 5 mM EGTA, pH 7.4) containing prostaglandin E1 (PGE1) (final concentration 1 µM). Following gentle mixing, residual white and red blood cells were removed by centrifugation at 100g for 15 min. The pellet was discarded and platelets in the supernatant were isolated from plasma via centrifugation at 800g for 15 min. The platelet pellet was washed twice with citrate dextrose saline buffer containing 10 mM sodium citrate, 150 mM sodium chloride, 1 mM EDTA, and 1% (w/v) dextrose (pH 7.4).

Sedimented platelets were resuspended in ice-cold 1:1 (v/v) mixture of platelet lysis buffer containing 0.5% Triton X-100, 30 mM HEPES, 150 mM NaCl, 2 mM EDTA, (pH 7.4), and Tyrode’s buffer. The lysis buffer also contained 2 µg/mL aprotinin, 10 µg/mL leupeptin, 10 mM sodium fluoride, 1 mM phenylmethylsulfonyl fluoride (PMSF), phosphatase inhibitor cocktail 2 (PIC2), phosphatase inhibitor cocktail 3 (PIC3), and 20 µM O-(2-Acetamido-2-deoxy-D-glucopyranosylidenamino) N-phenylcarbamate (PUGNAc). Disruption of the platelet membrane was achieved using ultrasonication in 15 s pulses (6x) at 70% maximum power. The lysed platelets were cooled on ice for one min between sonication cycles. Platelet lysate was separated from cellular debris by centrifugation at 1500 g. The protein concentration of the platelet lysate was estimated using bicinchoninic acid (BCA) assay (Thermo Scientific).

### Collagen affinity column preparation

Protein-derivitized agarose was generated using a modified version of a method reported previously by our group.^53^ Briefly, 500 µL of AminoLink^®^ Plus coupling resin (1 mL of a 50% slurry) was incubated with 5-fold diluted collagen (100 µg protein in 500 µL buffer (40 mM sodium citrate and 20 mM sodium carbonate, pH 10) at 37°C for 4 hrs on a rotisserie shaker. 500 mM sodium cyanoborohydride in 1X PBS was added (final concentration, 45.5 mM), and the sample was incubated at 37°C for 2 hrs with mixing. The resin was washed twice with 1X PBS, and incubated with 50 mM sodium cyanoborohydride and the sample was incubated again for 2 hrs at 37°C with mixing. The unreacted aldehyde sites remaining on the beads were blocked via incubation with 1 M Tris-HCl (pH 7.6) in the presence of 50 mM sodium cyanoborohydride. The sample was incubated at 37°C for 1 hr with mixing. The resin was washed 3X using 1M NaCl, and 10X with 1X PBS. The columns were stored in 1X PBS containing 0.02% NaN_3_ at 4°C before use. Coupling efficiency was determined by BCA assay comparing the flow-through to starting solution. For both collagen I and collagen IV, coupling was > 99% with nearly 100 µg collagen loaded in each column.

### Platelet protein collagen-affinity enrichment

Platelet lysate (1 mg) in PBS containing 50 mM MnCl^2^ was incubated in each collagen- affinity column for 4 hrs at 37°C on a rotisserie shaker. Unbound proteins were removed by sequential washes (10X using 1X PBS). Bound proteins were eluted with 1% TFA. The eluent was neutralized with 2 M ammonium bicarbonate and dried via speedvac. Proteins in the eluent were denatured at 37°C using 8 M urea in 1 M ammonium bicarbonate and the disulfide bonds were reduced with 5 mM dithiothreitol (DTT). Reduced cysteine residues were alkylated at 37°C by the addition of 10 mM iodoacetamide in the dark. The denatured, alkylated proteins were digested by the addition of Lys-C/trypsin mix (Promega) each in a 1/50 enzyme/protein w/w ratio. The sample was incubated at 37°C for 1 hr, and the solution was diluted two-fold with 50 mM ammonium bicarbonate to reduce the concentration of urea to 4 M. Trypsin was added in a 1/30 enzyme/protein w/w ratio, and the solution incubated for an additional 1 hr. The solution was diluted four-fold to a final urea concentration of 2 M, and an additional aliquot of trypsin was added, and the solution was incubated at 37°C overnight. Following digestion, the sample was dried and dissolved in 1% TFA. The pH of the solution was acidified to pH < 2 with 10% TFA, and the tryptic digest was desalted using C18 stage tips (3M Empore, Minneapolis, MN, USA). Residual detergent was removed from label-free samples using strong-cation exchange (SCX) tips (Glygen Corp., Columbia, MD). Peptides were labelled with 10-plex TMT reagents (Thermo Scientific) as described elsewhere.^54, 55^ TMT reagents were prepared by the addition of 40 µL dry acetonitrile to each channel (final concentration 20 µg/µL). Dried peptides (5 µg) were dissolved in 200 mM HEPES, pH 8.5, and 5 µL of the appropriate TMT reagent was added to each sample. TMT channels 126 and 127N were reserved for ‘blank’ channels representing the background noise arising from non-specific protein interactions with the blocked agarose beads. The remaining reagents were used to label samples in the order described in figure X. Labelling was performed at RT for 2 hrs and the reaction was quenched by the addition of 5% hydroxylamine and shaking for 30 min. Labelled samples were acidified by the addition of 1% TFA and pooled. The pooled samples were further acidified to pH > 2 by the addition of 10% TFA, and the pooled samples were desalted using C18 stage tips. Residual detergent was removed using SCX purification.

### Liquid-chromatography electrospray ionization tandem mass spectrometry

LC-MS experiments were performed using either a Q-Exactive mass spectrometer (Thermo Fisher Scientific) equipped with a Dionex Ultimate 3000 RSLC nano system with a 75 µm × 15 cm Acclaim PepMap100 separating column protected with a 2-cm guard column (Thermo Scientific). The mobile phase consisted of a 0.1% formic acid in water for solvent A and 0.1% formic acid in 95% aqueous acetonitrile for solvent B. The flow rate was 300 nL/min. TMT-labelled peptides were analyzed with an Orbitrap Fusion instrument (Thermo Scientific). Intact glycopeptide analysis was performed using either the Orbitrap Fusion or Orbitrap Lumos (Thermo Scientific) equipped with an EasyLC solvent pump. The gradient profile consisting of a stepwise gradient began with 4-7% B for 7 min, 7-40% B for 90 min, 40-90% B for 5 min, 90% B for 10 min, followed by equilibration (4% B) for 10 min. MS conditions were as follows, spray voltage was set to 2.2 kV, Orbitrap MS1 spectra (AGC 4 × 10^5^) were collected from 350-1800 m/z at an Orbitrap resolution of 120 K followed by data-dependent higher-energy collisional dissociation tandem mass spectrometry (HCD MS/MS) (resolution 60 K, collisional energy 35%, activation time 0.1 ms) of the 20 most abundant ions using an isolation width of 2.0 Da. Charge states were screened to reject both singly charged and unassigned ions; only charges between 2-6 were enabled. A dynamic exclusion window of 15 s was used to discriminate between previously selected ions.

### Database search

All TMT-labelled LC-MS/MS data were searched using Proteome Discoverer 2.2 (Thermo Scientific) searched against UniProtKB/Swiss-Prot human protein databases.

### Data processing software

Intact glycopeptide analysis was performed using GPQuest 3.0.^27^ All Venn diagrams were generated using eulerAPE software.^56^ Heat maps were generated using Gitools 2.3.1 software suite.

### Intact glycopeptide analysis

Raw MS files were converted to mzML format using MSConvert from Proteowizard and searched against peptide and glycan database using GPQuest.^27^ Oxonium ion containing spectra were extracted using HexNAc (204.087 Da) and at least one other oxonium ion (138.055 Da, 163.061 Da, 168.066 Da, 274.093 Da, 292.103 Da, and 366.140 Da) within a 5 ppm window as the selection criteria.

## Acknowledgements

This work was supported by Funding: National Cancer Institute, the Clinical Proteomic Tumor Analysis Consortium (CPTAC, U24CA210985), the Early Detection Research Network (EDRN, U01CA152813), National Heart Lung and Blood Institute, Programs of Excellence in Glycosciences (PEG, P01HL107153). The authors would like to thank Ms. Vicki Li for her assistance with sample preparation and helpful discussions.

## Author contributions

H.Z. and C.T. designed the study and wrote the manuscript; C.T. performed the experimental work; Y.H. and C.T. designed and performed the data analysis.

